# DNA N-gram Analysis Framework (DNAnamer): A generalized N-gram frequency analysis framework for the supervised classification of DNA sequences

**DOI:** 10.1101/2024.02.02.578674

**Authors:** John S. Malamon

**Author notes:** **CORRESPONDING AUTHOR:** John Stephen Malamon, PhD, University of Colorado, Anschutz Medical Campus, 1635 Aurora Court, Aurora CO 80045.

## Abstract

In 1948, Claude Shannon published a mathematical system describing the probabilistic relationships between the letters of a natural language and their subsequent order or syntax structure. By counting unique, reoccurring sequences of letters called N-grams, this language model was used to generate recognizable English sentences from N-gram frequency probability tables. More recently, N-gram analysis methodologies have been successful in addressing many complex problems in a variety of domains, from language processing to genomics. One such example is the common use of N-gram frequency patterns and supervised classification models to determine authorship and plagiarism. In this methodology, DNA is a language model where nucleotides are analogous to the letters of a word and nucleotide N-grams are analogous to the words of a sentence. Because DNA contains highly conserved and identifiable nucleotide sequence frequency patterns, this approach can be applied to a variety of classification and data reduction problems, such as identifying species based on unknown DNA segments. Other useful applications of this methodology include the identification of functional gene elements, sequence contamination, and sequencing artifacts. To this end, I present DNAnamer, a generalized and extensible methodological framework and analysis toolkit for the supervised classification of DNA sequences based on their N-gram frequency patterns.

## 1. Introduction

In 1948, Claude Shannon published “A Mathematical Theory of Communication.” [1], which provided a mathematical model defining the maximum information transmission rate and channel capacity that can be achieved given an arbitrarily small error probability. This groundbreaking work not only laid the foundation for modern telecommunication systems and information theory, but it also ignited the fields of stochastic modeling, data compression [2-4], cryptography [5, 6], and machine learning [7, 8]. This marked the birth of the Information Age. Moreover, Shannon arrived at the critical conclusion that communication signals and their information sources were statistically independent from the meaning of a given message. This finding is immensely important as it dispels the notion that more data implies more meaning. Rather, one must sift through layers of structured data and noise to extract the meaning.

In this seminal work, Shannon also developed a formal language model describing the probabilistic relationships between the letters in a natural language (English) and their subsequent order or syntax structure by counting reoccurring sequences of letters called N-grams. N*-*grams are unique, iterative sequences of *N* adjacent symbols. For example, the second-order N-gram set for the word “source” is [“so”, “ou”, “ur”, “rc”, “ce”]. These models are referred to as stochastic processes, as each N-gram within a set is represented as a random variable governed by a finite set of probabilistic states. Shannon’s remarkable ability to produce ordered or grammatically structured English sentences from N-gram probability frequency tables demonstrated that distinct and conserved syntax patterns and structures were embedded within natural languages. To this end, he stated, “It appears then that a sufficiently complex stochastic process will give a satisfactory representation of a discrete [information] source.”

The first and most fundamental hypothesis of this methodology, H_1_, is that the discrete information source of DNA (deoxyribonucleic acid), nucleotide sequences, can be adequately represented as a stochastic process similar to Shannon’s language model, where nucleotides [“A”,”C”,”G”,”T”] are analogous to the letters of a word and nucleotide sequence N-grams are analogous to the words of a sentence. This methodology also relies on two additional hypotheses. H_2_ requires that DNA N-grams such as bi-, tri-, and tetra-nucleotide pairings do not occur randomly, and H_3_ requires that DNA N-gram frequency patterns are conserved and identifiable. In fact, it has been demonstrated that di-, tri-, and poly-nucleotide repeats are not randomly distributed and are highly conserved with complex and distinct patterns [9-11]. Given H_2_ and H_3_, DNA N-gram frequency patterns can be leveraged to construct a generalized stochastic model that can be used to efficiently solve a wide variety of supervised classification and data reduction problems highly relevant to genomics and genetics.

Significant advancements have occurred in the past four decades in the fields of text categorization, authorship [12-17], and plagiarism detection [18, 19]. This progress has been achieved through the integration of novel N-gram analysis methodologies with supervised machine learning approaches. For example, anti-plagiarism tools based on N-gram analysis are commonplace and often positively regarded in educational institutions [20]. Since the sequencing of the human genome, numerous N-gram analysis methodologies have yielded an impressive range of inventive solutions by addressing relevant problems in genomics, genetics, and proteomics. These solutions include improving sequence alignment algorithms [21-23], describing microbial genetics [24, 25], estimating subcellular proteomes [26], classifying promotor sequences [27], and characterizing protein structures [28-30]. To this end, I present DNAnamer, or DNA N-gram Analysis Framework, a generalized DNA N-gram frequency analysis methodology and toolkit for the binary and multi-class supervised classification of DNA sequences.

## 2. Materials and methods

### 2.1. Data source and description

The human reference genome (hg38) and four additional reference genomes were randomly selected from the *Class Mammalia* and analyzed to demonstrate a specific use case and highlight the broad applicability of this methodology. Each species considered required at least 1,000 megabases (Mb) of nucleotide sequences, or roughly one-third of the human genome. The species used in these *in silico* experiments were *Homo sapiens* (human), *Miniopterus natalensis* (bat), *Elephas maximus* (elephant), *Phascolarctos cinereus* (koala), and *Delphinus delphis* (dolphin). The five reference genomes were downloaded from the National Center for Biotechnology Information’s genome resource database using the ‘biomartr’ package [31]. The reference genomes totaled approximately 3.3 (human), 1.8 (bat), 3.4 (elephant), 3.2 (koala), and 2.5 (dolphin) billion nucleotides.

### 2.2. DNA N-grams and frequency tables

DNA N*-*grams are unique, iterative sets of adjacent nucleotide sequences. The order of an N-gram is equivalent to the number of adjacent nucleotide sequences. For example, “AA” represents a second-order N-gram. The total number of DNA N-gram sequences in a set is equal to 4^*O*^, where *O* is the order or the number of consecutive nucleotides. For example, the second-order DNA N-gram nucleotide sequence set is [“AA”,”AC”,”AG”,”AT”,”CA”,”CC”,”CG”,”CT”,”GA”,”GC”,”GG”,”GT”,”TA”,”TC”,”TG”, “TT”].

Because this study involved analyzing nucleotide frequency patterns, N-gram frequency tables were calculated for all reference genomes using the ‘seqinR’ package [32]. The analysis of variance (ANOVA) test [33] was used to calculate the difference in N-gram frequency means within and across species. **Equation 1** demonstrates a second-order N-gram frequency (*f*) matrix. The ‘*no*.*tests’* was equal to the total DNA sequence length divided by the segment length. The ‘*no*.*tests’* was selected to provide at least 200 validation samples per experiment, which provided an ample range of frequency distributions and a minimum performance evaluation resolution of 0.5%. N-gram frequency tables were calculated for all randomly selected training and validation DNA segment sets and transposed into a matrix where the columns were equal to the total cumulative number of N-grams, and the rows were equal to the ‘*no*.*tests’* for a given experiment. The maximum cumulative number of N-grams is given in **Equation 2**.

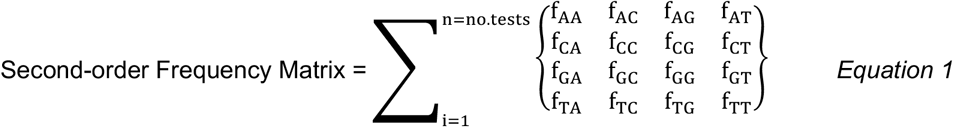

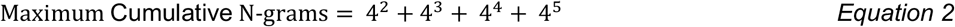

### 2.3. Randomness analysis

This methodology relies on the assumption that DNA N-gram frequencies are not randomly distributed. Specifically, there are two relevant dimensions, or definitions of randomness implied. First, in-species randomness was tested to ensure non-randomness across the set of all N-grams within each species. Second, cross-species randomness was tested to ensure nonrandomness between two given species. Two well-accepted randomness methodologies were applied to directly test these assumptions. First, the Wald-Wolfowitz runs test [34] was used to test the two randomness hypotheses. This approach makes minimal assumptions regarding the underlying N-gram frequency distributions. Second, Bartels’ test for randomness [35] was applied. This test is a ranked version of von Neumann’s ratio test for randomness [36]. In summary, von Neumann’s approach evaluates statistical trends in the successive means of a number sequence. Bartels’ ranked test offers several distinct advantages over von Neumann’s ratio test.

### 2.4. Experimental design

A graphic illustration of this methodology and the *in silico* experimental design was provided in **Figure 1**, where sets of DNA segments ranging from 10 to 100kb in length were randomly selected from each of the five species’ (human, bat, elephant, koala, and dolphin) reference genomes to develop training and validation sets, each consisting of 20Mb of continuous DNA sequence. The main objectives of this experiment were to eliminate the aforementioned assumptions, identify N-gram frequency patterns, and classify unknown contiguous DNA sequences based on established patterns. This experiment also examined the relationship between this model’s classification performance as a function of N-gram order and the DNA segment length. Therefore, second, third, fourth, and fifth-order N-grams were analyzed and compared. Relatively small DNA segments up to 100kb in length were analyzed to discover the upper limits of this model’s classification performance. Specifically, unknown human DNA segments totaling 20Mb and ranging from 10 to 100kb were classified against each of the four other mammals. The five sequence segment lengths tested were 10, 20, 40, 80, and 100kb. This yielded a total of eighty binary classification experiments. The number of validation tests for each experiment was equal to the total sequence length (20Mb) divided by the segment length. For example, the 100kb segment length yielded 200 validation sequences.

**Figure 1.**
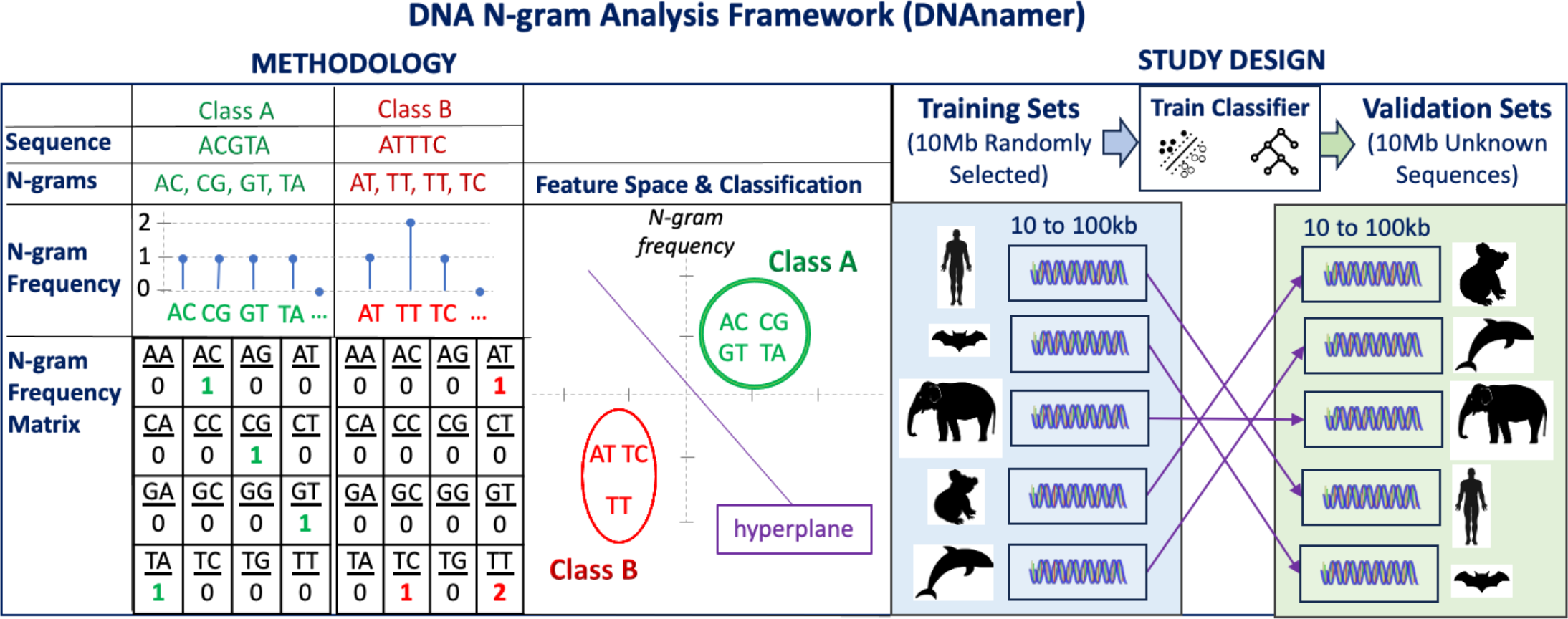
Graphic abstract of the methodology and experimental design. The DNAnamer (DNA N-gram Analysis Framework) methodology is illustrated to the left, where two simple examples of second-order DNA N-gram sequences are provided. N-gram frequency tables were created, and DNA sequences were classified based on their N-gram frequency patterns. The Support Vector Machine was used for binary classification and the random Forest Classifier was used to analyze more than two classes. The experimental study design is depicted to the right, where a range of DNA segments ranging from 10 to 100kb in length were randomly selected from each of the five species’ (human, bat, elephant, koala, and dolphin) reference genomes to develop training sets (blue panel). Validation sets of unknown DNA segments ranging from 1 to 100kb in length were classified for all five species (light green panel).

#### 2.4.1. Model training

DNA sequences were randomly sampled across the entire reference genome of each species to create balanced training sets each consisting of 20Mb of continuous sequence. N-gram frequency distribution priors were calculated using only the training sets. The validation sets (20Mb) consisted of continuous DNA segment sets ranging from 10 to 100kb and were used to classify unknown DNA sequence segments from all five organisms.

#### 2.4.2. Binary classification

The Support Vector Machine (SVM) methodology has proven to be a highly versatile and effective classification approach well-suited for a variety of biomedical and genomic applications [37-40]. Importantly, the SVM has been successfully employed to identify species based on their DNA sequences, a technique also known as DNA barcoding [41]. For this reason, the SVM methodology [42] was selected for the binary classification experiments. All eighty experiments were performed using the same SVM parameters. Namely, a radial kernel was selected with a cost of 3 and a sigma value of 0.2. 50% to 50% sample hold-out validation was used to calculate the area under the receiver operating characteristic curve, or AUC. The AUC was provided as a function of the N-gram order (second, third, fourth, and fifth) and DNA segment length (10-100kb).

#### 2.4.3. Multi-class classification

The Random Forest (RF) methodology has also proven to be a highly versatile and effective classification approach for a variety of biomedical and genomic applications [43-46]. RFs have also been successfully employed to perform DNA barcoding [47-49]. For this reason, the Random Forest classification methodology [50] was selected for multi-class classification. All experiments were performed under the same SVM parameters. Namely, a maximum of 300 trees was specified with 100 permutations and 40 mtry. The mtry parameter specifies the maximum number of input features or variables available to a decision tree. Specifically, this methodology was used to measure relative variable importance and classify unknown DNA segments using models trained with DNA from all five organisms. 10Mb of sequence from each species was used to develop these training and validation sets. The AUC was provided as a function of the N-gram order and DNA sequence segment length, yielding twenty multi-classification experiments.

#### 2.4.4. Benchmark Analysis

Finally, computational performance benchmarking was performed on the human reference genome. Total runtimes (seconds), the number of iterations per second (iterations/second), the total memory allocated (GB), and core seconds (runtime/cores) were recorded on a 2.4 GHz 8-Core Intel Core i9 processor with 64GB of available memory. Performance benchmarks were provided for all necessary functions and the five DNA segment lengths. 20Mb of DNA sequences were processed for each of the eight benchmark experiments. Fifth-order N-gram analysis was performed to provide a realistic estimation of the computational resources required to replicate these experiments. All analyses were performed using the R statistical language version 4.3.2 [51].

## 3. Results

### 3.1. DNAnamer

The DNA N-gram Analysis Framework (DNAnamer) is a novel N-gram frequency analysis framework for the supervised classification of DNA sequences and is available as an R software package or library. Documentation and vignettes with detailed code demonstrations are available at https://github.com/jmal0403/DNAnamer/wiki. All major classification experiments performed herein can be reproduced using the vignettes and sample code. In summary, DNAnamer provides a highly generalized, efficient, and extensible analytical framework that can be readily applied to take on a variety of classification and data reduction problems in the fields of genomics and genetics.

### 3.2. DNA N-gram frequencies are nonrandom and conserved

The Wald-Wolfowitz runs test and Bartels’ ranked test confirmed in-species and cross-species N-gram frequency nonrandomness. These two randomness tests were repeated for the five N-gram orders and across the five species, yielding p-values less than 2.2e-16 in all instances. Thus, N-grams were far from randomly distributed within and across species. **Figure 2A-D** provided box plots of the second-order N-gram frequencies for 200 randomly selected DNA segments 100kb in length calculated for human versus bat, elephant, koala, and dolphin reference genomes. The ANOVA test revealed many statistically significant differences in the N-gram frequency means between and among species (**Supplemental Figure 1**), further supporting the nonrandomness hypothesis. Many statistically significant (p-value<0.05, **Supplemental Table 1**) cross-species N-gram frequency patterns were observed, providing a wealth of independent random variables for classification. The cross-species N-gram frequency correlation matrix was given in **Figure 2E**. N-gram frequency patterns were highly conserved across the five species, all yielding correlation coefficients greater than 0.95. Thus, H_2_ and H_3_ have been directly tested to reveal many unique and identifiable N-gram frequency patterns in and across species.

**Figure 2.**
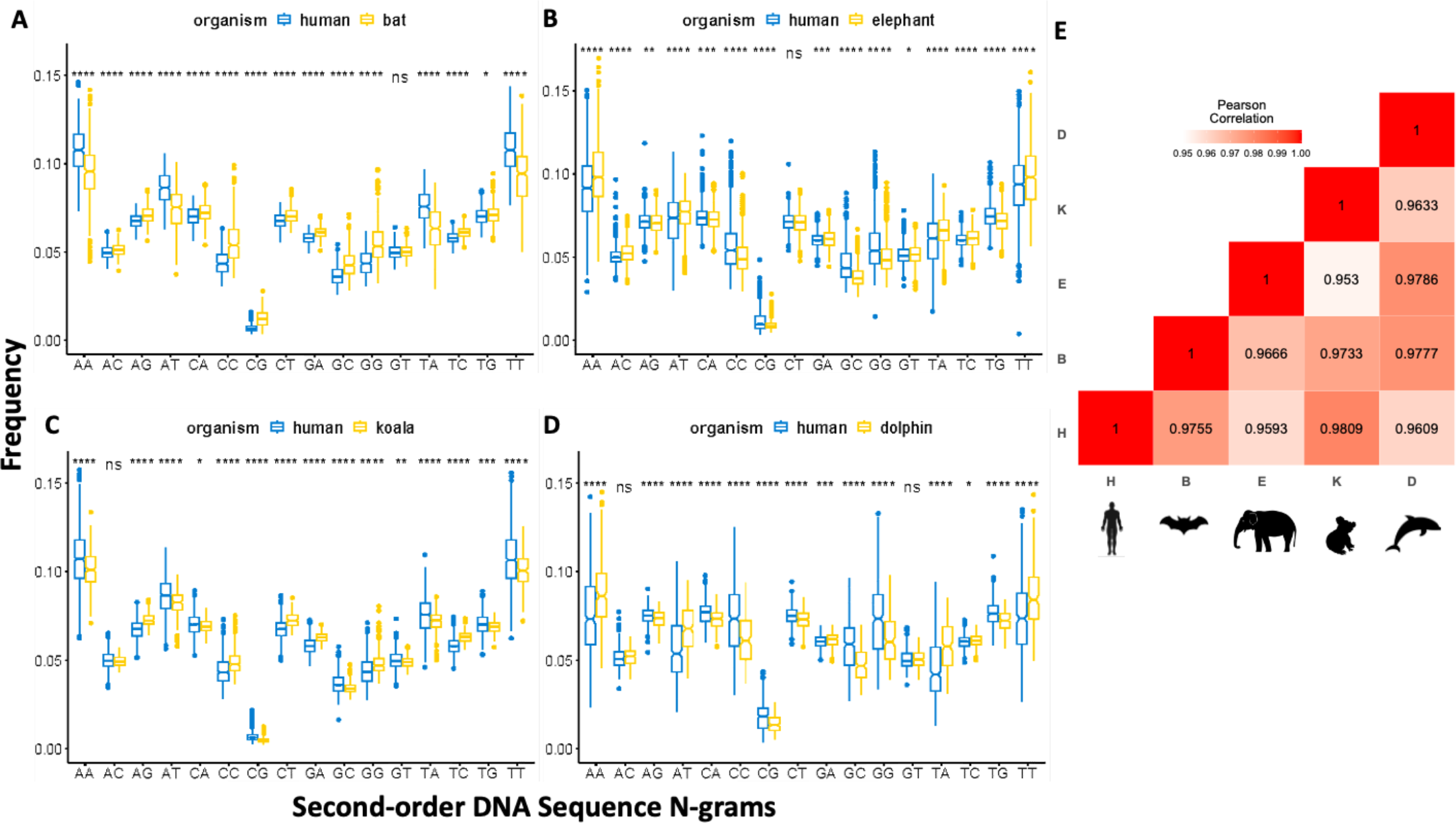
Second-order DNA N-gram frequencies for human versus bat, elephant, koala and dolphin. Using training sets of 20MB and DNA segment lengths of 100kb, second-order N-gram frequencies were calculated for A) human versus bat; B) human versus elephant; C) human versus koala; and D) human versus dolphin. The ANOVA was used to measure the differences in the group means between the two species. The following p-value notation was used: *p > 0*.*05 [12], p < 0*.*05 (****\*****), p < 0*.*01(****\*\*****), p < 0*.*001 (***), ns = not significant*. Panel E provided a Pearson’s cross-correlation matrix for the second-order N-gram frequencies comparing all five species.

### 3.3. N-gram variable importance using Random Forests

**Supplemental Table 2** provided the top 40 most important N-grams for all five species, sorted by their importance in humans. In summary, 36 out of the 40 top N-grams were fifth-order, and the remaining four were fourth-order. Although second- and third-order N-grams were important in the RF model, fourth- and fifth-order N-grams provided additional statistical discrimination and cross-species variability. Interestingly, variable importance varied significantly among the five species, demonstrating that N-gram frequency patterns were both unique and conserved.

### 3.4. Classification performance is a positively correlated with N-gram order and segment length

**Figure 3A** provided the AUCs grouped by N-gram order and as a function of the DNA segment length for unknown human sequence segments versus the other four species. **Figure 3C** provided a table containing a summary of the number of tests performed for each binary classification experiment. In summary, even low-order N-gram analysis performed well in binary classification mode, with second-order analysis averaging AUCs of 0.851, 0.916, 0.919, 0.949, and 0.951 for 10, 20, 40, 80, and 100kb DNA segments, respectively. Fifth-order analysis averaged AUCs of 0.973, 0.993, 0.993, 0.999, and 1, respectively. In binary mode, all 200 test sequences were correctly classified in the human genome versus the four other mammalian genomes. For reference, 100kb is approximately 1/33,000, or 0.000003%, of the size of the human genome. **Figure 4** provided the AUCs grouped by the N-gram order and as a function of the segment length used when classifying all five species. The contingency tables for each multi-class experiment were provided in **Supplemental Table 3**. Fifth-order classification AUCs ranged from 0.85 to 0.979 and increased with the DNA segment length.

**Figure 3.**
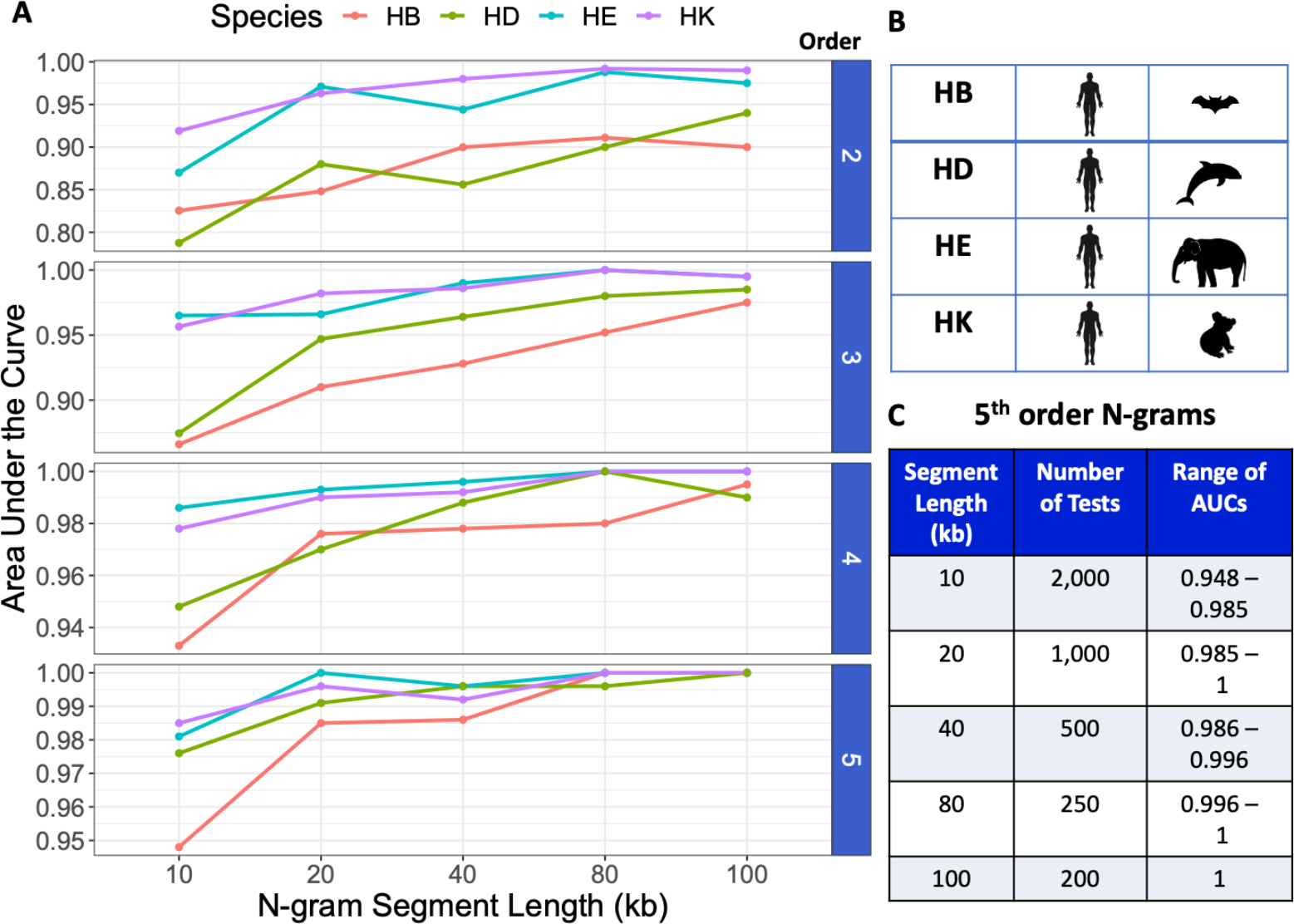
Supervised binary classification using the Support Vector Machine. Panel A provided the area under the receiver operating characteristic curve or AUC as a function of DNA N-gram segment length and grouped by the N-gram order. Each plot in Panel A was scaled independently. Model performance was assessed at each DNA sequence segment length (10, 20, 40, 80, and 100kb) and for each order (2, 3, 4, 5). This experiment was repeated for humans versus the four other species, bat (HB), elephant (HE), koala (HK), and dolphin (HD) as depicted in Panel B. Panel C provided the number of validation experiments performed and the AUC range at each DNA segment length for fifth-order N-gram analysis.

**Figure 4.**
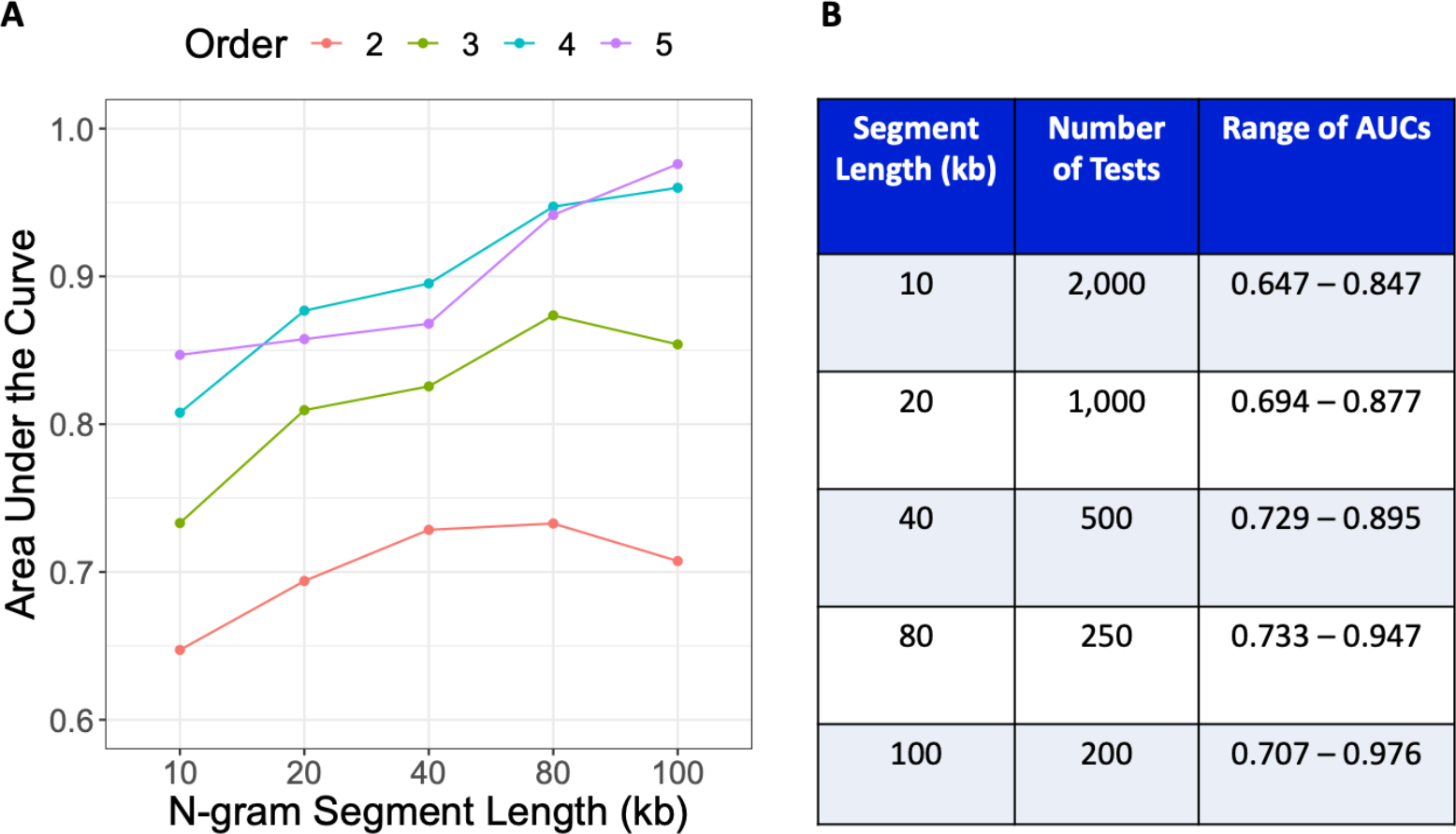
Supervised multi-class classification using Random Forests. Panel A provided the area under the receiver operating characteristic curve or AUC as a function of the DNA N-gram segment length and were colored by the N-gram order. Model performance was assessed at each DNA sequence segment length (10, 20, 40, 80, and 100kB) and for each order (2, 3, 4, 5). Panel B provided the number of validation experiments performed and the AUC range at each DNA segment length.

### 3.5. Computational performance benchmarking

Fifth-order N-grams were used to provide an approximation of this model’s computational performance using a total of 20Mb of DNA sequence for each benchmark. Three functions were required to perform supervised classification. They were *getSequence(), getFreqDF()*, and *binaryClassification()*. The *getSequence()* function randomly selected and constructed the contiguous DNA segments. On average, this function required 19 seconds to process 20MB of DNA using 4.7GB of memory. The *getFreqDF()* function calculated the N-gram frequency tables and required between 30 and 70 seconds to complete while using 12 to 32GB of memory. This function’s runtime and memory increased with the number of tests, as expected with a larger matrix. Finally, the *binaryClassification()* function performed SVM classification and required 1 to 44 seconds to process using less than 1.4GB of memory. The average combined processing time for a fifth-order binary classification experiment was only 90 seconds, or 12 seconds per processing core.

## 4. Discussion

DNANAMER provides an efficient, accurate, and extensible methodology and analysis toolkit designed to tackle a broad range of complex and computationally expensive classification problems applicable to the biomedical sciences, genomics, and genetics. This methodology achieves high levels of classification accuracy using minimal information. This innovative toolkit for stocastic modeling and machine learning has been validated, documented, and is available as a complete and well-tested R package.

To establish theoretical feasibility, a number of fundamental methodological assumptions and hypotheses have been addressed. Specifically, it has been demonstrated that DNA N-grams are highly conserved and exhibit distinct and identifiable patterns, making them excellent for classification. Next, a series of carefully crafted DNA barcoding experiments were designed and executed to offer a concrete example and application for this innovative approach. In completing these classification experiments, it was demonstrated that model performance consistently increased with N-gram order and DNA segment length. Thus, it would seem logical to conclude that classification accuracy could improve by applying higher-order N-gram analysis.

Like all models, this one has both strengths and limitations. One such limitation of this toolkit is that it used 32GB of memory to calculate the largest N-gram frequency table. This isn’t ideal, but it can easily run on a modern laptop computer. The small sample size of the species used in this DNA barcoding demonstration was a further limitation of this study. Although hundreds of millions of DNA sequences were efficiently analyzed and accurately classified, it will be interesting to see how this approach performs with organisms that exhibit higher levels of sequence variability and toxonomic distance.

Because the five species analyzed in this study have relatively high sequence homology, more genetically divergent species may exhibit even more distinct and identifiable N-gram frequency patterns. Among this model’s main strengths are efficiency, accuracy, and generality. Efficiency allows for data reduction applications and the analysis of higher-order N-grams. Most importantly, the generality of this methodology makes it ideal for a wide array of problems that we face in the ever-growing and dynamic field of genomics. In an era of ‘big data’, it is critical to remember that more data does not necessarily imply more meaning.

Broader application and integration was the driving force behind designing, building, and validating this methodology and analysis toolkit. As such, I will conclude with three specific and promising applications for this methodology. First and foremost, this approach is ideal for classifying functional genetic elements. As previously outlined, N-gram analysis methodologies coupled with machine learning approaches have been highly successful in identifying functional gene elements such as promotor regions and genomic islands. Expanding on these approaches for identifying novel functional gene elements would greatly increase our knowledge of genetics and could improve our understanding of human disease. A second common and practical application for this methodology is the identification of cross-species DNA contamination. Bacterial DNA contamination occurs in DNA extraction kits, PCR (polymerase chain reaction) reagents, and model organisms. This methodology could be fashioned to detect bacterial and other DNA contaminants along with sequencing artifacts caused by non-biological processes. Finally, this approach also allows for generative modeling through the simulation of more biologically accurate *in silico* DNA sequences. These simulated sequences can be used to advance *in silico* spike-in procedures to improve structural variant, copy number variation, and retrotransposon detection and quality control tools.

## Data Availability

All data used to conduct this study can easily be downloaded from the National Center for Biotechnology Information’s genome resource database.

## Acknowledgment

The author has no acknowledgments.

---

**Table 1.** Computational performance benchmarks for fifth-order N-gram analysis. Total Runtime (seconds), Core Seconds, Maximum Memory (GB) and Iterations per second are provided. Benchmarking was performed for all necessary processing steps for fifth-order N-gram analysis, with segments ranging from 10 to 100kb and totaling 20MB of DNA sequence. Core Seconds were equal to the Total Runtime divided by the number of processing cores (N=8).

| ***Function*** | **Segment Length (kb)** | **Total Runtime (seconds)** | **Core Seconds** | **Maximum Memory (GB)** | **Iterations per second** |
| --- | --- | --- | --- | --- | --- |
| *getSequence* | 10 | 19.2 | 2.40 | 4.75 | 0.052 |
| *getSequence* | 20 | 18.7 | 2.34 | 4.77 | 0.054 |
| *getSequence* | 40 | 18.8 | 2.35 | 4.74 | 0.053 |
| *getSequence* | 80 | 18.5 | 2.31 | 4.77 | 0.054 |
| *getSequence* | 100 | 18.9 | 2.36 | 4.76 | 0.053 |
| *getFreqDF* | 10 | 69.6 | 8.70 | 32.3 | 0.014 |
| *getFreqDF* | 20 | 45.4 | 5.68 | 16.7 | 0.022 |
| *getFreqDF* | 40 | 34.4 | 4.30 | 12.7 | 0.029 |
| *getFreqDF* | 80 | 32.6 | 4.08 | 11.7 | 0.031 |
| *getFreqDF* | 100 | 30.8 | 3.85 | 11.5 | 0.033 |
| *binaryClassification* | 10 | 44.2 | 5.53 | 1.4 | 0.022 |
| *binaryClassification* | 20 | 10.4 | 1.30 | 0.74 | 0.096 |
| *binaryClassification* | 40 | 2.57 | 0.32 | 0.39 | 0.389 |
| *binaryClassification* | 80 | 0.89 | 0.11 | 0.22 | 0.112 |
| *binaryClassification* | 100 | 0.71 | 0.09 | 0.19 | 0.14 |

