## Supplement for "DNA N-gram Analysis Framework (DNAnamer): A generalized N-gram frequency analysis framework for the supervised classification of DNA sequences"

This appendix has been provided by the authors to give readers additional information about the work.

**Supplementary appendix for:**

***DNA N-gram Analysis Framework {DNA-NAMER}: A Generalized N-gram Frequency Analysis Framework for the Supervised Classification of DNA Sequences***

### Contents

**Supplemental Table X**. . ........................................................................................................................ X
**Supplemental Table X**. . ........................................................................................................................ X

**Figure 1**. **Second-order DNA N-gram Frequencies for Five Species.**

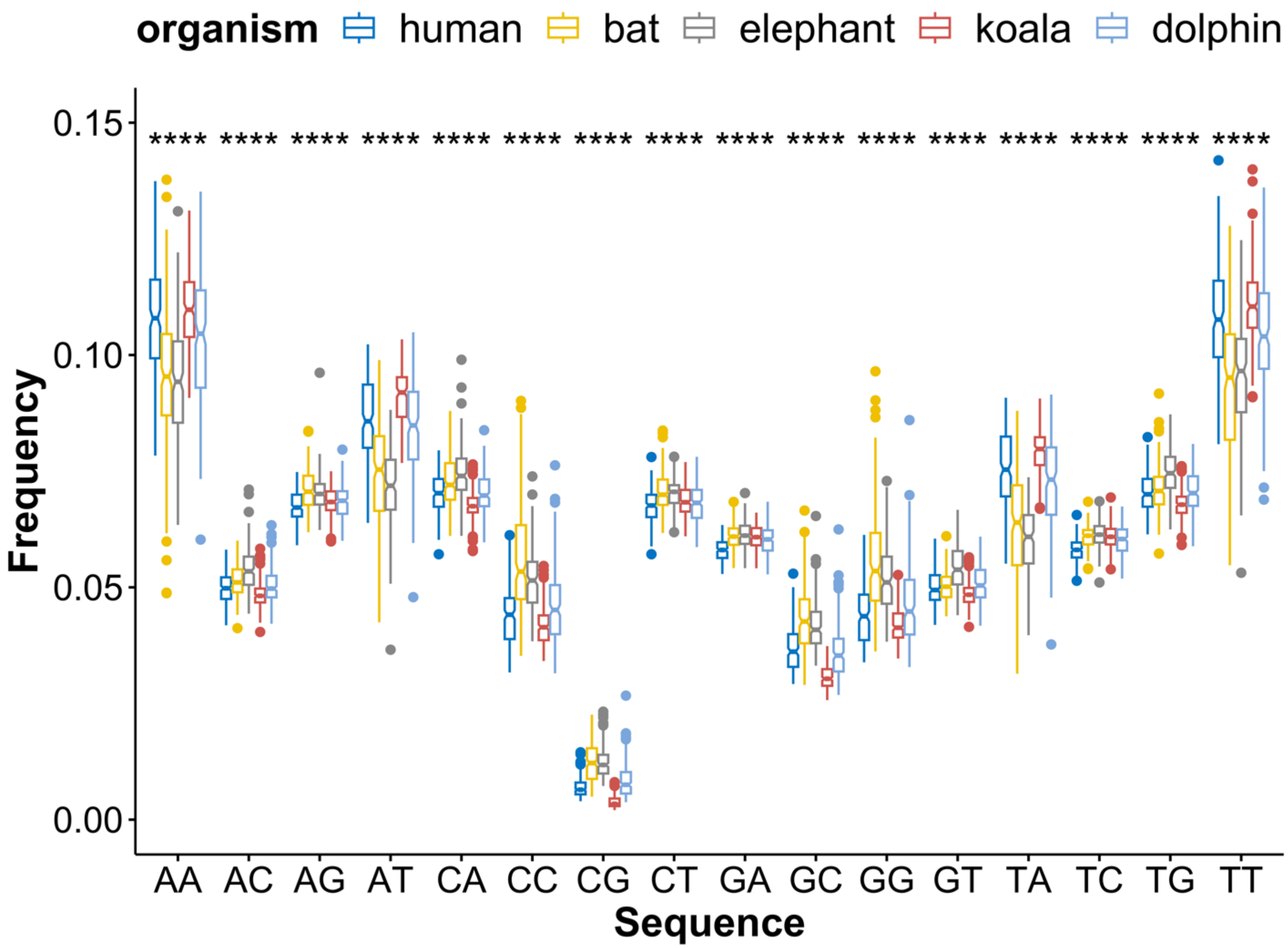

**Supplemental Table 1.** ANOVA Results for Second-order DNA N-gram sequences. The Bonferroni methodology was used to calculate 'p-adj'.

| | ***Human versus Bat*** | | | ***Human versus Elephant*** | | |
|---|---|---|---|---|---|---|
| ***N-gram*** | **p-value** | **p-adj** | **Sig.** | **p-value** | **p-adj** | **Sig.** |
| *AA* | 1.21E-93 | 1.94E-92 | **** | 1.25E-21 | 2.00E-20 | **** |
| *AC* | 3.40E-25 | 5.44E-24 | **** | 9.84E-15 | 1.57E-13 | **** |
| *AG* | 5.65E-68 | 9.04E-67 | **** | 2.29E-03 | 3.66E-02 | * |
| *AT* | 7.82E-189 | 1.25E-187 | **** | 8.39E-14 | 1.34E-12 | **** |
| *CA* | 2.45E-46 | 3.92E-45 | **** | 1.30E-06 | 2.08E-05 | **** |
| *CC* | 5.08E-151 | 8.13E-150 | **** | 5.49E-26 | 8.78E-25 | **** |
| *CG* | 4.10E-166 | 6.56E-165 | **** | 9.67E-26 | 1.55E-24 | **** |
| *CT* | 1.04E-73 | 1.66E-72 | **** | 8.61E-03 | 1.38E-01 | ns |
| *GA* | 1.23E-41 | 1.97E-40 | **** | 1.49E-04 | 2.38E-03 | ** |
| *GC* | 6.63E-140 | 1.06E-138 | **** | 4.22E-77 | 6.75E-76 | **** |
| *GG* | 5.65E-137 | 9.04E-136 | **** | 4.85E-30 | 7.76E-29 | **** |
| *GT* | 9.16E-01 | 1.00E+00 | ns | 2.27E-01 | 1.00E+00 | ns |
| *TA* | 5.13E-195 | 8.21E-194 | **** | 4.19E-28 | 6.70E-27 | **** |
| *TC* | 3.19E-61 | 5.10E-60 | **** | 8.02E-06 | 1.28E-04 | *** |
| *TG* | 1.79E-06 | 2.86E-05 | **** | 4.23E-26 | 6.77E-25 | **** |
| *TT* | 9.07E-131 | 1.45E-129 | **** | 2.07E-09 | 3.31E-08 | **** |
| | ***Human versus Koala*** | | | ***Human versus Dolphin*** | | |
| *AA* | 9.19E-13 | 1.47E-11 | **** | 2.39E-11 | 3.82E-10 | **** |
| *AC* | 8.51E-02 | 1.00E+00 | ns | 2.46E-01 | 1.00E+00 | ns |
| *AG* | 6.05E-46 | 9.68E-45 | **** | 7.41E-05 | 1.19E-03 | ** |
| *AT* | 2.14E-12 | 3.42E-11 | **** | 4.49E-18 | 7.18E-17 | **** |
| *CA* | 3.99E-03 | 6.38E-02 | ns | 1.32E-12 | 2.11E-11 | **** |
| *CC* | 2.32E-18 | 3.71E-17 | **** | 3.41E-16 | 5.46E-15 | **** |
| *CG* | 3.00E-32 | 4.80E-31 | **** | 4.17E-13 | 6.67E-12 | **** |
| *CT* | 5.72E-45 | 9.15E-44 | **** | 4.19E-06 | 6.70E-05 | **** |
| *GA* | 8.47E-68 | 1.36E-66 | **** | 2.41E-03 | 3.86E-02 | * |
| *GC* | 1.98E-15 | 3.17E-14 | **** | 1.84E-27 | 2.94E-26 | **** |
| *GG* | 6.79E-16 | 1.09E-14 | **** | 8.28E-16 | 1.32E-14 | **** |
| *GT* | 1.92E-04 | 3.07E-03 | ** | 9.72E-01 | 1.00E+00 | ns |
| *TA* | 8.67E-12 | 1.39E-10 | **** | 8.54E-22 | 1.37E-20 | **** |
| *TC* | 1.36E-59 | 2.18E-58 | **** | 6.32E-02 | 1.00E+00 | ns |
| *TG* | 1.90E-07 | 3.04E-06 | **** | 2.12E-14 | 3.39E-13 | **** |
| *TT* | 9.43E-15 | 1.51E-13 | **** | 2.39E-11 | 3.82E-10 | **** |

**Supplemental Table 2.** Top 40 Most Important N-grams for Classifying Human DNA using Random Forests.

| *N-gram* | Human | Bat | Dolphin | Elephant | Koala |
|---|---|---|---|---|---|
| *GCTA* | 5.3615 | 3.2779 | 5.4778 | 2.2553 | 3.2965 |
| *CATG* | 5.0365 | 3.2528 | 2.2059 | 0.1865 | 2.4685 |
| *TCTCA* | 4.2708 | 2.1922 | 2.7316 | 1.8844 | 0.8692 |
| *TCGAT* | 4.1077 | 5.21 | 3.1231 | 4.516 | 3.5103 |
| *ATCGA* | 3.9004 | 5.4768 | 4.6215 | 4.976 | 4.3406 |
| *TGCA* | 3.7099 | -0.2346 | 2.7907 | 2.1967 | 4.2637 |
| *CTCTC* | 3.667 | 4.4578 | 1.3065 | 2.6237 | 1.2283 |
| *GAGAG* | 3.4757 | 4.5807 | 2.6773 | 2.3394 | 1.8128 |
| *TAGC* | 3.4589 | 3.3071 | 2.4019 | 1.5525 | 2.3465 |
| *GTCGA* | 3.3512 | 0.0986 | 3.0612 | 3.7658 | 2.1298 |
| *TCGAC* | 3.2708 | 1.3889 | 1.9881 | 4.6277 | 2.3776 |
| *AGTCG* | 3.1944 | 2.2251 | 2.7695 | 4.3304 | 2.5590 |
| *TCCGG* | 3.1919 | 1.716 | 3.6209 | 0.1544 | 2.0995 |
| *AGTCA* | 3.1817 | 2.2979 | 2.8608 | 0.9569 | 3.5970 |
| *GGTTA* | 3.1518 | 2.4437 | 2.0341 | 3.2765 | -1.0017 |
| *CTGCA* | 3.104 | -0.5165 | 0.8839 | 2.1208 | 2.9461 |
| *ATGGT* | 3.1019 | 2.8716 | 1.4910 | 0.977 | 2.1187 |
| *GGACC* | 3.0988 | 0.6354 | 2.2730 | 1.0017 | 0.5777 |
| *GTCTC* | 3.0956 | 1.3281 | 0.9931 | 1.1968 | 1.9609 |
| *CATCG* | 3.0916 | 2.0126 | 1.1517 | 2.07 | 1.4541 |
| *AGCTA* | 2.9584 | 2.4362 | 3.136 | 3.136 | 3.0215 |
| *GTCCG* | 2.9294 | 1.0107 | 1.7446 | 1.7446 | 1.3972 |
| *TAGCT* | 2.8935 | 0.473 | 2.1545 | 2.1545 | 2.3348 |
| *GATAG* | 2.7881 | -0.9011 | 2.2433 | 2.2433 | 2.7547 |
| *ACGCG* | 2.7677 | -0.7968 | 1.2157 | 1.2157 | 1.9561 |
| *TAACC* | 2.758 | 1.3974 | 4.0261 | 4.0261 | 1.0017 |
| *ACCAT* | 2.7186 | 2.3253 | 2.005 | 2.005 | -1.0017 |
| *CATAG* | 2.6693 | 2.5844 | 2.7818 | 2.7818 | 1.419 |
| *GGTCC* | 2.6592 | 0.6911 | 1.8397 | 1.8397 | -1.0017 |
| *CATGA* | 2.6066 | 0.55 | -0.0681 | -0.0681 | -0.8213 |
| *AGAGA* | 2.5894 | 3.4093 | 2.0012 | 2.0012 | 1.6271 |
| *CAGTT* | 2.5669 | 0.1714 | 2.731 | 2.731 | 0.933 |
| *GCATG* | 2.559 | 2.1274 | 2.6106 | 2.6106 | 1.5979 |
| *TCTCT* | 2.549 | 2.8475 | 1.6678 | 1.6678 | 1.3886 |
| *TGCAG* | 2.5357 | 0.7599 | 0.1243 | 0.1243 | 2.4749 |
| *TGACT* | 2.5191 | 1.4502 | 1.4335 | 1.4335 | 3.1648 |
| *TCGA* | 2.5008 | 2.8362 | 3.4861 | 3.4861 | 2.237 |
| *AGTC* | 2.4889 | 2.537 | 3.9762 | 3.9762 | 1.9512 |
| *ACTAG* | 2.4795 | 2.5321 | 0.2344 | 0.2344 | 2.6885 |
| *TGCAA* | 2.4705 | 1.3696 | 0.5186 | 0.5186 | 1.6595 |

**Supplemental Table 3.** Contingency Tables for Multi-class Classification using the Random Forest Methodology with fifth-order N-grams and DNA segments length.

| | | | ***Segment Length = 10kb*** | | | |
|---|---|---|---|---|---|---|
| ***Species*** | **Human** | **Dolphin** | **Elephant** | **Human** | **Koala** | **Class Error** |
| ***Bat*** | 64 | 22 | 3 | 10 | 1 | 0.36 |
| ***Dolphin*** | 21 | 62 | 10 | 7 | 0 | 0.38 |
| ***Elephant*** | 4 | 10 | 86 | 0 | 0 | 0.14 |
| ***Human*** | 17 | 9 | 1 | 69 | 4 | 0.31 |
| ***Koala*** | 0 | 4 | 0 | 1 | 95 | 0.05 |
| | | | ***Segment Length = 20kb*** | | | |
| ***Bat*** | 71 | 9 | 1 | 13 | 6 | 0.29 |
| ***Dolphin*** | 5 | 83 | 6 | 6 | 0 | 0.17 |
| ***Elephant*** | 4 | 6 | 88 | 2 | 0 | 0.12 |
| ***Human*** | 8 | 4 | 2 | 85 | 1 | 0.15 |
| ***Koala*** | 1 | 4 | 0 | 0 | 95 | 0.05 |
| | | | ***Segment Length = 40kb*** | | | |
| ***Bat*** | 85 | 7 | 0 | 6 | 2 | 0.15 |
| ***Dolphin*** | 2 | 95 | 0 | 3 | 0 | 0.05 |
| ***Elephant*** | 0 | 0 | 100 | 0 | 0 | 0 |
| ***Human*** | 2 | 1 | 0 | 97 | 0 | 0.03 |
| ***Koala*** | 0 | 0 | 0 | 0 | 100 | 0 |
| | | | ***Segment Length = 80kb*** | | | |
| ***Bat*** | 85 | 7 | 0 | 6 | 2 | 0.15 |
| ***Dolphin*** | 2 | 95 | 0 | 3 | 0 | 0.05 |
| ***Elephant*** | 0 | 0 | 100 | 0 | 0 | 0 |
| ***Human*** | 2 | 1 | 0 | 97 | 0 | 0.03 |
| ***Koala*** | 0 | 0 | 0 | 0 | 100 | 0 |
| | | | ***Segment Length = 100kb*** | | | |
| ***Bat*** | 92 | 4 | 3 | 1 | 0 | 0.08 |
| ***Dolphin*** | 0 | 99 | 1 | 0 | 0 | 0.01 |
| ***Elephant*** | 2 | 0 | 98 | 0 | 0 | 0.02 |
| ***Human*** | 0 | 0 | 0 | 96 | 4 | 0.04 |
| ***Koala*** | 0 | 0 | 1 | 0 | 99 | 0.01 |
